# Seasonality and competition select for variable germination behavior in perennials

**DOI:** 10.1101/2022.01.13.476161

**Authors:** Hanna ten Brink, Thomas R. Haaland, Øystein H. Opedal

## Abstract

The common occurrence of within-population variation in germination behavior and associated traits such as seed size has long fascinated evolutionary ecologists. In annuals, unpredictable environments are known to select for bet-hedging strategies causing variation in dormancy duration and germination strategies. Variation in germination timing and associated traits is also commonly observed in perennials, and often tracks gradients of environmental predictability. Although bet-hedging is thought to occur less frequently in long-lived organisms, these observations suggest a role of bet-hedging strategies in perennials occupying unpredictable environments. We use complementary analytical and evolutionary simulation models of within-individual variation in germination behavior in seasonal environments to show how bet-hedging interacts with fluctuating selection, life-history traits, and competitive asymmetries among germination strategies. We reveal substantial scope for bet-hedging to produce variation in germination behavior in long-lived plants, when ‘false starts’ to the growing season results in either competitive advantages or increased mortality risk for alternative germination strategies. Additionally, we find that lowering adult survival may, in contrast to classical bet-hedging theory, result in less spreading of germination by decreasing density-dependent competition. These models extend insights from bet-hedging theory to perennials and explore how competitive communities may be affected by ongoing changes in climate and seasonality patterns.

## Introduction

Accurate timing of seasonal phenology is key to population persistence in unpredictable environments. The timing of emergence can have strong and direct effects on individual fitness, and is expected to be subject to strong selection driving local adaptation (Donohue et al. 2010). Indeed, emergence behavior (e.g. patterns of germination or hatching) is often found to vary in predictable ways along environmental gradients (Meyer et al. 1995; Venable 2007; Wagmann et al. 2012; Simons 2014; Pinceel et al. 2017; Rubio de Casas et al. 2017; Torres-Martínez et al. 2017; Scholl et al. 2020). In most plants and many animals, the timing of emergence is controlled by dormancy (Vleeshouwers et al. 1995; Finch-Savage and Leubner-Metzger 2006; Baskin and Baskin 2014), and the evolution of emergence behavior is thus tightly linked to the evolution of dormancy mechanisms (Varpe 2017).

Theoretical models of seed dormancy and germination behavior have a long history (Cohen 1966; Venable and Lawlor 1980; Ellner 1985; Geritz et al. 2018; Hughes 2018; Kortessis and Chesson 2019). Most of these, however, have focused on specific systems such as annual plants in desert environments. This system provides a natural starting point because one striking observation demands an explanation: a fraction of the seeds produced each year fail to germinate the next year, and instead lie dormant for another year before germinating at the beginning of the second growing season following their dispersal (Gremer and Venable 2014). While this reduction in number of seedlings may seem a waste of resources most years, such a strategy has been identified as a risk-spreading adaptation to avoid complete recruitment failure in the event of a bad year. Prolonged seed dormancy and variation in germination timing have come to represent the archetype of a bet-hedging strategy, i.e. a genotype-level strategy that sacrifices some short-term (arithmetic-mean) fitness gains in order to lower its fitness variance over time (Levins 1962; Cohen 1966; Venable 2007). Such bet-hedging strategies have received considerable empirical and theoretical attention as a major mode of adaptation to unpredictable environments (Seger and Brockmann 1987; Simons 2011; Starrfelt and Kokko 2012).

Empirical and theoretical studies of bet-hedging have yielded a good understanding of how different types of risk-spreading adaptations co-evolve, interact, and cancel each other out. For example, if phenotypic polymorphisms or continuous variation ensures that fitness correlations among related individuals are sufficiently low, then there is less need for other costly bet-hedging strategies at the individual level such as ‘safer’ offspring phenotypes that are weaker competitors but better able to cope with environmental variation (Venable and Brown 1988; Starrfelt and Kokko 2012; Haaland et al. 2020; Escobar et al. 2021). The scope for bet-hedging is also lowered if the spatiotemporal scale of environmental variation (what is often called environmental ‘grain’ in bet-hedging terms) already ensures that related individuals experience different conditions and thus uncorrelated fitness returns. In a spatially fine-grained environment relatives may be exposed to a wide range of different conditions at any given time (as opposed to coarse-grained environments where all individuals experience the same conditions), and in such cases, genotype-level diversification or other bet-hedging strategies are not selectively favored (Levins 1962; Starrfelt and Kokko 2012). Furthermore, longer lifespans with a larger number of selective events over which fitness can accumulate (e.g. an individual experiencing multiple consecutive breeding seasons with variable conditions) also reduces the need for any variance reduction if fitness payoffs among selective events are uncorrelated (Haaland et al. 2019).

Given these insights, it is unsurprising that most models of bet-hedging, as well as many of the best-documented empirical examples, consider rather specific types of short-lived organisms with discrete generations occupying highly variable, coarse-grained environments, such as germination strategies of desert annuals, or overwintering strategies for organisms living in ephemeral ponds (Furness et al. 2015; García-Roger et al. 2016; Wang and Rogers 2018). However, this focus has also led to a knowledge gap regarding the evolution of seemingly ‘risk-spreading’ traits in longer-lived organisms, as these may require other explanations than bet-hedging in its strictest sense. Variation in seed germination behavior is seen not only in annuals, and is at least sometimes associated with variation in seed size (Rees 1996; Simons and Johnston 2000; Susko and Lovett-Doust 2000; Norden et al. 2009; Harel et al. 2011; Martins et al. 2019). A classic example of within-individual variation in seed size is seed heteromorphism, as observed e.g. in many Asteraceae and Chenopodiaceae (reviewed in Venable 1985; Imbert 2002), where distinct seed morphs differ in their dispersal abilities and germination behavior (Venable et al. 1987; Brändel 2004; Fumanal et al. 2007; Yao et al. 2010; Wang et al. 2012). Continuous variation in seed size is also common in natural populations, and may relate to variation in germination behavior (Pélabon et al. 2021). Within species, larger seeds are generally more likely to germinate and/or germinate earlier than do smaller ones (Biere 1991; Simons and Johnston 2000; Tremayne and Richards 2000; Galloway 2001; Pélabon et al. 2005), but the opposite pattern of smaller seeds breaking dormancy and germinating earlier is also observed (Martins et al. 2019).

Seed size variation is thus a common mechanism for producing seeds with variable duration of dormancy and/or variable germination time both within individuals as well as within an inflorescence (Pélabon et al. 2021). As argued, existing bet-hedging models alone are not able to explain the ubiquity of seed heteromorphism or continuous seed size variation as observed in perennials (Martins et al. 2019). Although several other non-mutually exclusive hypotheses exist, including but not limited to variance-sensitivity due to asymmetrical fitness costs versus benefits (Bednekoff 1996; Bårdsen et al. 2008), fluctuating selection or fluctuations in priority effects (Chesson 2000; ten Brink et al. 2020), competitive asymmetries among offspring types (Geritz 1995) and negative frequency-dependent selection (Metcalf et al. 2015; Poethke et al. 2016), these have seldom been analyzed jointly (but see Rees et al. 2004). In Scholl et al.’s (2020) recent analysis of the flora of the south-east United States, annuals were not detectably more likely to exhibit seed heteromorphism than were perennials, suggesting that selection for within- and/or among-individual variation in germination behavior may be common also in long-lived plants where bet-hedging is less strongly favored.

Here, we explore the evolution of individual variation in dormancy of long-lived plants in seasonally varying environments, using both analytical and simulation models covering a range of ecological scenarios. First, our analytical model derives general predictions for when within- and/or among-plant variation in germination behavior can be favored as an adaptation to seasonal environmental variation in perennials. To this end we use an adaptive dynamics approach assuming that plants produce two discrete seed morphs, an ‘early’ seed with a short dormancy period, and a ‘late’ seed with a long dormancy period (for example, ‘small’ and ‘large’ seeds, respectively). This analysis examines the conditions favoring plants producing seeds of variable dormancy duration (frequencies of seed morphs different from 0 or 1), and identifies the shape of the function relating the (rainfall-dependent) benefits and costs of producing early relative to late seeds as the key determinant of whether such mixed strategies are stable. The potential for among-individual variation (presence of evolutionary branching points) is also examined. We complement this analytical model with individual-based evolutionary simulations of dormancy duration modelled as a near-continuous trait via freely-evolving weekly germination probabilities. This model thus examines the predictions from the analytical model without the restriction of only allowing two discrete seed morphs, and makes explicit the mechanisms affecting benefits and costs of early and late seeds. Specifically, early seeds germinating during the dry season gain a competitive advantage if they survive until reproduction, but suffer high mortality if there is not enough rain between their germination and the beginning of the wet season (‘false starts’). We let spatiotemporal rainfall patterns vary predictably and unpredictably within and among years, thus emulating the wet- and dry-season dynamics seen in large parts of the tropics where intermittent dry-season rains can trigger germination of certain seed types. However, we note that this setup applies equally well to other stochastic seasonal changes between harsh and favorable conditions, such as the onset of warmer spring weather in temperate regions where ‘false starts’ of warm temperatures may still be followed by harmful frosts. In concert, our two models allow us to tease apart the relative effects of bet-hedging, competition and fluctuating selection on the evolution of variable germination strategies.

### 1. Selection for within-plant variation in germination behavior

We first present an analytical model where we assume that plants can produce two seed morphs, here ‘early’ and ‘late’ seeds, that may differ in e.g. size or shell thickness and hence the amount of rainfall required for germinating (Norden et al. 2009). Each year consists of a dry and a wet season, where the dry season is characterized by scarce intermittent rainfalls, affecting the germination and survival probability of early seeds. These early seedlings emerging during intermittent dry-season rains have a high probability of dying due to drought, and therefore only a small chance to recruit into the adult populations, but may persist if there is sufficient excess rain in the following weeks leading up to the wet season. Late seeds, on the other hand, remain dormant through any intermittent rains, and germinate at the beginning of the safe wet season. However, any early seedlings that survived until the beginning of the wet season may now have a competitive advantage over late seeds, as they have a head start to growth, and therefore a higher probability of recruitment into the adult population. Thus, the relative advantage *b* of being an early *vs*. a late seed in a given year depends on the amount of excess dry-season rainfall in that year, a random environmental variable we term *X*.

We designate as *f* the fraction of early seeds produced by a plant, so that a fraction 1–*f* are late seeds. We model a density-regulated population of long-lived plants, where adult plants survive with probability *s*_a_, and all seeds compete for recruiting into the vacant space 1–*s*_a_ in the adult population. We incorporate asymmetric competition among early and late seeds in a ‘lottery’ for vacant space. Early seeds get the relative weight *b*(*x*)*f* against the weight 1–*f* of all late seeds, such that the probability of drawing an early seed in the lottery for vacant space becomes

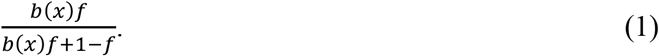

Next, assuming that the population is sufficiently isolated (no immigration) and that the quality of environmental conditions *X* vary randomly from year to year without any autocorrelation, the relative fitness *w*(*f’, f*) of strategy *f’* (a rare mutant) against a resident population that has strategy *f* is given by the geometric average growth rate, which is the expectation over the log growth rates *λ*, i.e.

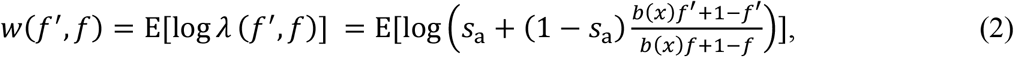

where E indicates the expectation over all environmental conditions *X*=*x* (see appendix A for derivation of *λ*). The sign of *w*(*f’,f*) determines whether the mutant can invade the population (positive sign) or not (negative sign) – examples are shown in Fig. 1a-b where expectations are calculated over 10.000 realizations of *x*∼Normal(0,1).

**Figure 1:**
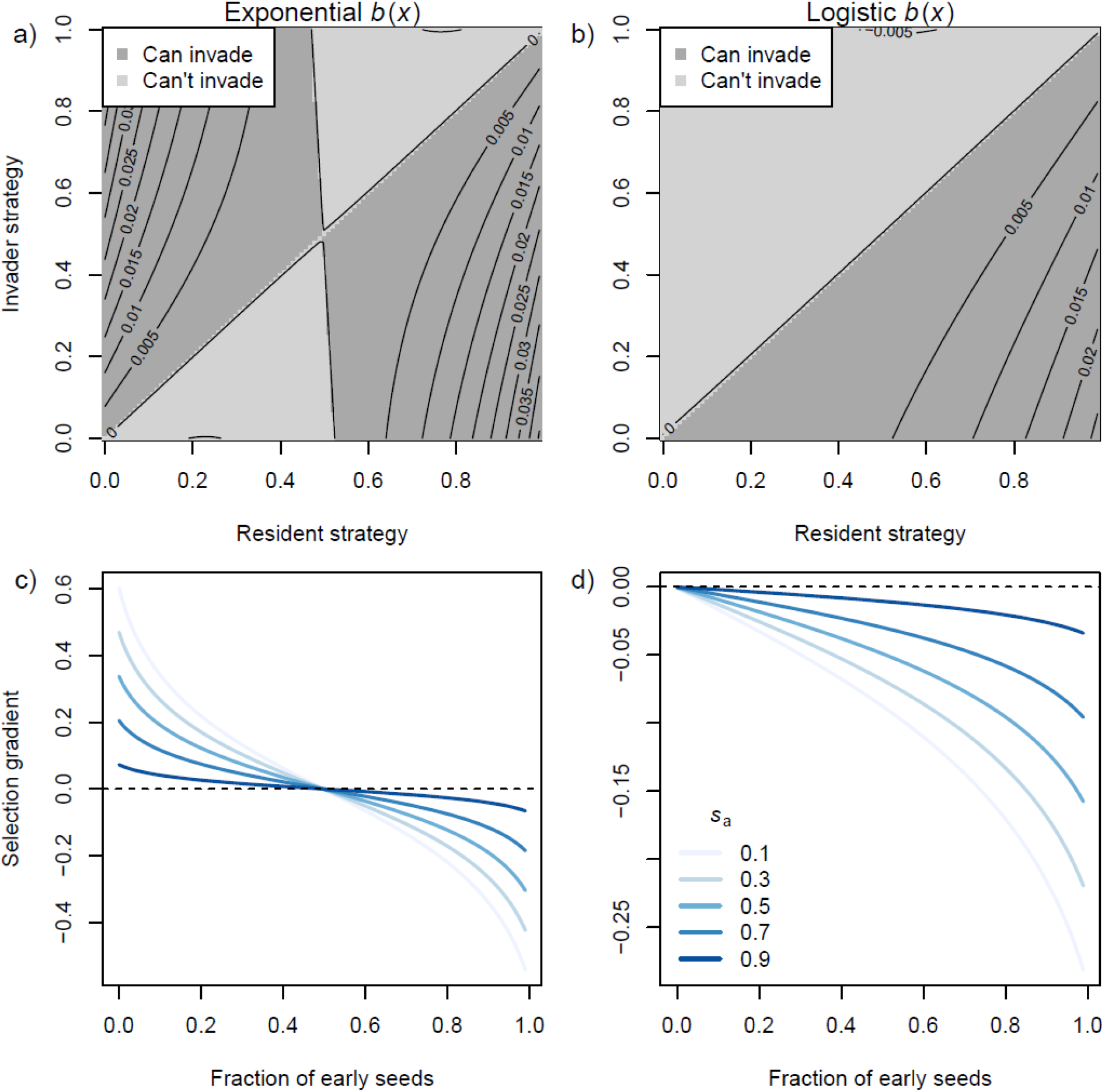
Results from adaptive dynamics model of the evolution of variable seed germination timing (strategy *f* defined as fraction of early-germinating seeds) in a long-lived plant (*s*_a_=0.9). Results are shown when the sum of the benefits and costs of early relative to late germination relate to environmental conditions *x* (excess dry-season rainfall) with an exponential (left column, using *β*=1) or logistic (right column, using *α*=1) function *b*(*x*). Top row: Pairwise invasibility plots showing fitness surfaces of invasion fitness (eq. 2) as a function of the resident strategy *f* and the invader strategy *f’*. Dark shaded regions indicate regions where the mutant strategy can invade. Bottom row: Selection gradients (eq. 4) on *f’* showing the direction and strength of evolution for different *f* (x-axis) and values of adult survival *s*_a_ (line color). Values of *f* where the lines cross 0 (horizontal dashed line) represent convergence stable ESSs (see eqs. 5&6).

We now use an adaptive dynamics approach to analyse the evolutionary consequences of equation 2 (Kisdi and Meszéna 1995; Geritz et al. 1998). Assuming that E[*X*]=0 (the expectation for ‘excess’ rain is zero) and that the two seed types fare equally well in average years when there is neither an excess nor deficit of rain between the early seeds germinating and the end of the dry season (i.e., *b*(0)=1), we can examine the conditions leading to an advantage for early seeds.

The selection gradient acting on a rare mutant with a germination strategy (fraction of early seeds) *f’* slightly different from the resident population’s strategy *f*, is obtained by differentiating *w*(*f’, f*) with respect to *f’* and evaluating at *f’*=*f*. Differentiating eq. 2 yields:

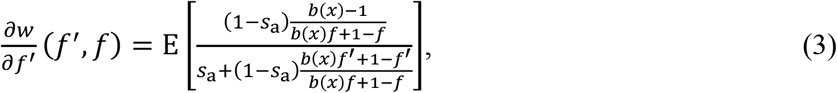

such that the selection gradient becomes

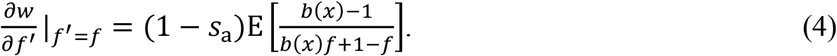

Intuitively, the strength of selection decreases with increasing adult survival 0<*s*_a_<1, as the competition among seedlings for annual recruitment contributes a smaller part of the population. Examples of selection gradients for two functions *b*(*x*) with different *s*_a_ are shown in Fig. 1c-d. Cases where 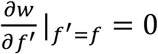 (colored lines cross the dashed black lines in Fig. 1c-d) represent so-called singular strategies (Geritz et al. 1998). In order to determine whether these are attractors or repellors we can examine the sign of the derivative of eq. 4 taken with respect to the resident strategy *f*:

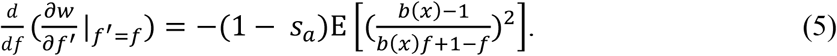

Because the squared term is positive, eq. 5 is always negative, indicating that all singular strategies are attractors. Further, computing the second derivative of equation 2 with respect to the mutant strategy *f*’ and again evaluating at *f’*=*f* allows us to determine whether the singular strategies are evolutionarily stable strategies (negative sign) or branching points (positive sign). Thus, the ESS criterion is:

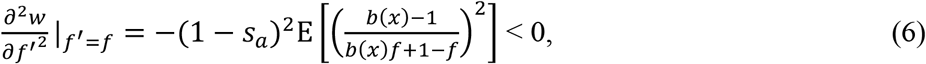

which reveals already that 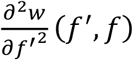 is always negative (as both squared terms are always positive), thus there is no potential for evolutionary branching and any singular strategy is necessarily a convergence stable ESS (Geritz et al. 1998). In other words, in this simple case of two discrete seed morphs (early and late seeds), the evolution of their frequency *f* cannot reach two distinct values at evolutionary equilibrium.

The need to compute expectations over realizations of the random variable *X* makes it difficult to exactly identify ESSs in the current model. However, we can approximate selection gradients using the delta method if we assume that *X* follows a sufficiently simple distribution with all odd-numbered moments equal to 0 (E[*X*]=0, symmetric distribution), a small variance *V*, and even-numbered moments have an upper bound depending on *V*. Then, recalling that *b*(0)=1 by assumption, we obtain the approximation

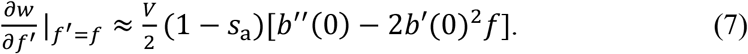

Setting this selection gradient equal to 0, we see that there can be one singular strategy only,

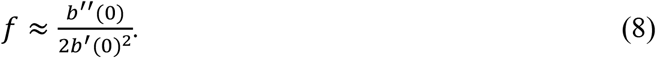

In order for such an ESS to exist and represent a mixed strategy, it much lie between 0 and 1, with conditions given by the shape of the function *b*(*x*), which determines the advantage of early relative to late seeds for a given level of excess dry-season rain *x*. Firstly, to ensure *f*>0, *b*’’(0) > 0 is demanded (eq. 8), i.e. *b*(*x*) must be accelerating (convex) around 0. Secondly, to ensure *f*<1, eq. 8 gives the condition *b’’*(0) < 2*b’*(0)^2^, i.e. it must not accelerate too fast (there is an upper bound to how convex it can be, given by the steepness of the function). For example (as shown in Fig. 1a,c), if *b*(*x*) = e^-*βx*^, then we have an ESS for all *β*>0, since *b*’’(0) > 0 and *b’’*(0) < 2*b’*(0) is fulfilled. The ESS becomes *f* = *b’’*(0) / 2*b’*(0) = 0.5. In contrast, if *b* is a logistic function *b*(*x*)=2/[1+e^-*αx*^] (Fig. 1b,d), the conditions for an ESS are not fulfilled: *b’*(0) = 0 and *b’’*(0)<0 whenever *α*>0.

Overall, this model reveals that there is ample scope for within-individual variation in germination strategies when plants can evolve to produce fractions *f* and 1-*f* of distinct early and late seeds, but that among-individual variation in *f* should not occur in such a scenario because all ESSs are convergence stable. The evolution of a mixed strategy 0 < *f* < 1 requires accelerating rewards for germinating early with increasing amounts of excess rainfall *x*, but only within a limited range. If rewards are not accelerating, then *f* evolves towards 0 (if benefits of being early are not sufficient, there is no point in producing risky early-germinating seeds), whereas if rewards accelerate too much, *f* evolves towards 1 and all seeds germinate early. Interpreting *b*(*x*) as the sum of the functions relating early seed dry-season survival as a function of *x* and the competitive advantage of early seeds (given their survival) over late-germinating seeds in the lottery for vacant space in the adult population, this analysis provides a baseline expectation for our individual-based model with continuous variation in germination time.

### 2. Individual-based model of mean and variance in dormancy Model setup

Here, we build an individual-based simulation model that allows plants to produce a continuous distribution of seeds (rather than just e.g. ‘early’ and ‘late’ seeds). We model a population of asexually reproducing perennials occupying a wet-dry seasonal tropical environment where the wet and dry seasons both last half the year (26 weeks). We assume that seed germination is determined by the dormancy period *d*, which functions as an obligate ‘biological clock*’* preventing germination until a certain time *d* has passed (see table 1 for an overview of all mathematical notation and baseline parameter values used in the individual-based model). Germination of a seed occurs with probability 1 as soon as its age (in weeks) is greater than *d*_z_. We set no constraints on the evolution of flexible dormancy strategies, as probability of germination is determined by 27 haploid genes *d*_*i*_, with *i* between 1 and 27, each corresponding to a week in the dry half of the year, and the 27^th^ indicating germination at the start of the wet season. Each gene determines the probability of a seed germinating in the matching week, and therefore has values bounded by 0 and 1, with the condition that the sum of all genes always equals 1. The actual phenotypic trait value (dormancy period *d*_z_) of an offspring *j* is randomly drawn from the probability distribution determined by its parent’s genetic values.

**Table 1:** Overview of variables and parameters of the individual-based model.

| Parameter in text | Description | Default values and range |
| --- | --- | --- |
| **A) Abiotic environment parameters** |  |  |
| $m$ | Mode of the beta distribution determining weekly dry-season rainfall (see eq. 9) | Default 0.5, we explore [0, 1] |
| $v$ | Variance of the beta distribution | $\frac{1}{36} \frac{m+1}{2-m}$ |
| $R_t$ | Realized rainfall in a particular week $t$. | [0, 1]. |
| **B) Survival function parameters** |  |  |
| $a$ | Age of seedling. | |
| $S_{\min}$ | Lower asymptote of the seedling survival function (survival probability given $a=1$ and $R_t=0$). | Default 0.6. |
| $S_{\max}$ | Upper asymptote of the seedling survival function (survival probability when $a$ and $R_t$ are large). | Default 0.997. |
| $s_{\text{ad}}$ | Weekly survival probability of plants older than 26 weeks | Default 0.997, we also examine $s_{\text{ad}} = 0.993$ |
| $\delta_1$ | Affects the steepness of the survival function. | Default 0.2. |
| $\delta_2$ | Enhances the effect of rain on the survival function. | Default 0.8. |
| $a_{\text{inf}}$ | Age (weeks) at which the survival function increases most steeply. | Default 5. We also examine $a_{\text{inf}} = \{4, 7\}$. |
| **C) Genes and trait values** |  |  |
| $d_i$ | Genetic values determining weekly probabilities of breaking dormancy, $i \in \{1, 2, \dots, 27\}$. | [0, 1]. Require $\sum_i d_i = 1$. |
| $d_z$ | Realized individual dormancy period (in weeks). | |
| **D) Demographic and evolutionary parameters** |  |  |
| $K$ | Carrying capacity. | Default 5000. |
| $\gamma$ | Yearly fecundity per adult plant. | Default 50, we also examine $\gamma = 130$ |
| $\mu$ | Per-locus mutation rate. | Default 0.1. |
| $\mu\sigma$ | Size of mutational effects (The standard deviation of the Gaussian distribution from which the new trait value is chosen). | Default 0.001. |
| $N_0$ | Starting population size. | Default equal to $K$. |
| $c$ | Strength of competitive asymmetries (competitive advantage to germinating earlier), see eq. 11. | Default 1. We also examine $c = \{0.5, 1.2\}$. |

#### Population initialization

Each replicate simulation starts with *N*_0_ adult individuals. Each of these individuals are independently assigned a random combination of gene values, where starting *d*_*i*_ values are uniformly distributed between 0 and 1. After assigning each gene a random value, we standardize all values such that the sum of an individual’s 27 genes equals 1. We start the simulations with ample standing genetic variation for computational purposes.

#### Abiotic conditions

We model time steps of one week and assign the first 26 weeks of a year as the dry season and the last 26 weeks as the wet season. Dry-season rainfall varies stochastically within years, with 26 weekly rainfalls randomly drawn at the start of the year from a beta distribution bounded between 0 (no rain) and 1 (high rain). Weekly rainfall in the wet season is always equal to 1. We parameterize the beta distribution in terms of its mode *m*, such that the shape parameters are given by

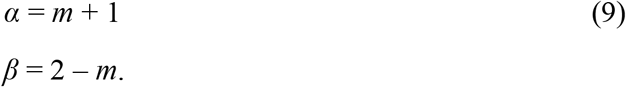

The mode mainly affects the average rainfall (the higher the wetter), but note that it also affects weekly rainfall variance within years 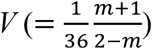, which is highest when *m*=0.5.

To tease apart effects of bet-hedging resulting from the spatiotemporal scale of environmental stochasticity, we examine evolutionary outcomes under coarse-grained and fine-grained rainfall variation. In coarse-grained environments, we assume that all seeds experience the same sequence of weekly dry-season rainfalls ***R*** (a vector of 26 values) but ***R*** differs among years. Under fine-grained environmental variation, different ***R*** vectors are assigned to individual seeds within years at random. Specifically, we produce 1000 sequences of weekly rainfall upfront and assign each seed a random ***R*** chosen among these. Thus, experienced rainfall will vary a lot among individuals within a year in this scenario, but average rainfall will vary less across years. In both scenarios, the weekly rainfall in the wet season (week 27– 52) is constant and equal to 1, which represents high rainfall.

#### Survival of seedlings and adults

After setting the weekly rainfalls of a particular year, we determine the survival probability of seedlings and adults in that year. We assume that survival increases both with rainfall and seedling age. The probability of survival for an individual *i* with age *a*_*i,t*_ in a given week *t* with rainfall *R*_*t*_ (or *R*_*i,t*_ in the fine-grained scenario) equals

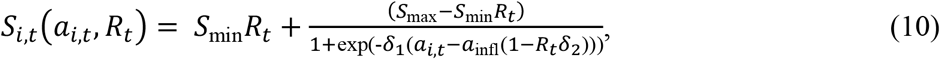

i.e. a sigmoid function of seedling age, the lower asymptote of which increases with increasing rainfall (Fig. 2A, B). Note that rainfall also affects the horizontal position of the function (how far ‘along’ the x-axis the seed has come in terms of survival), and that the strength of this effect increases with the parameter *δ*_2_.

**Fig. 2:**
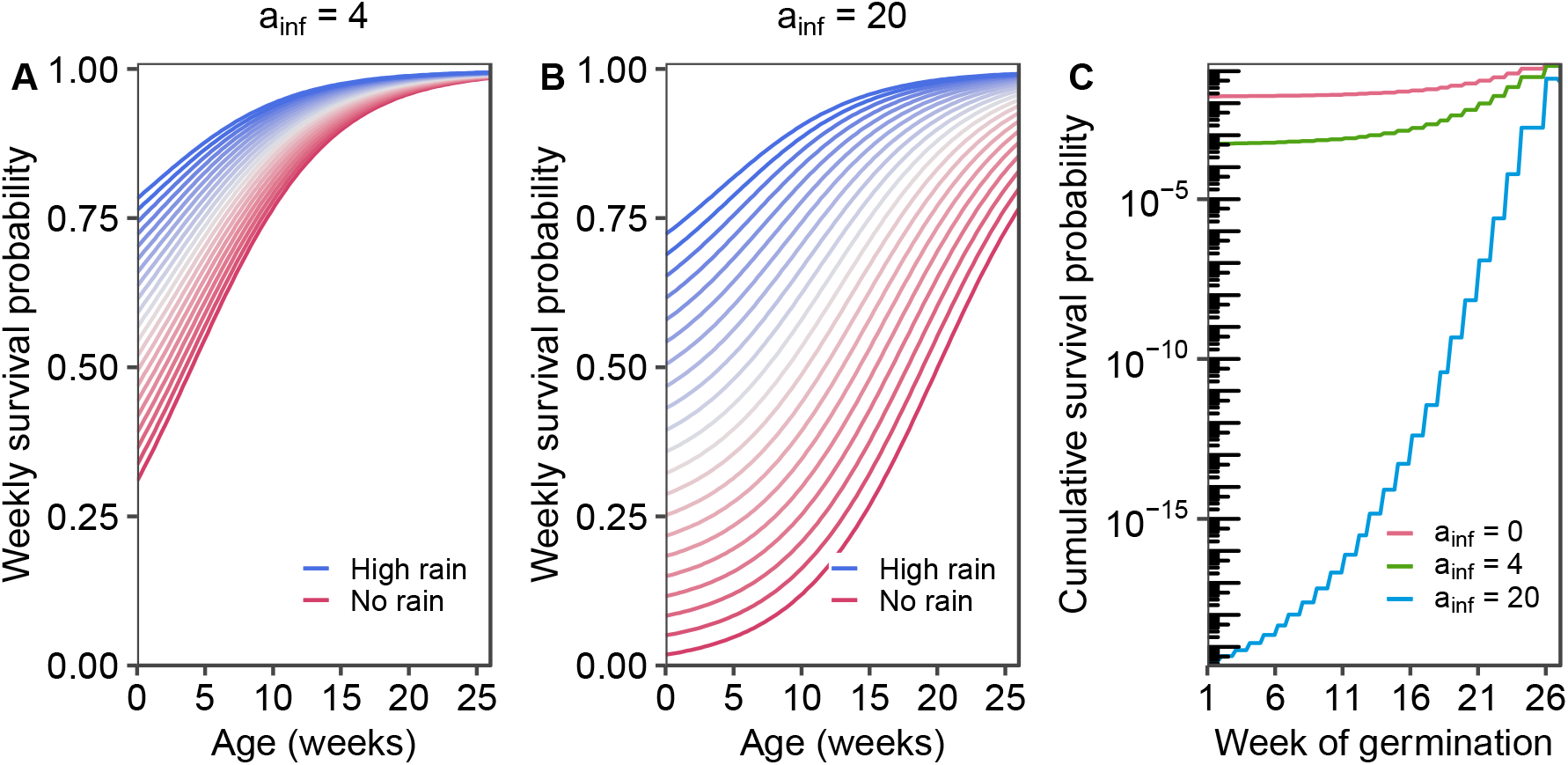
Seedling survival functions. A and B: Weekly survival probability for different amounts of rainfall (blue lines: ***R***=1; red lines: ***R***=0) and varying the inflection point *a*_inf_ of the juvenile survival function *S*_*i,t*_ (panels). C: Effect of week of germination (x-axis) and *a*_inf_ (colored lines) on the probability of a seedling surviving its first year (52 weeks since germination), assuming a worst-case scenario with no rainfall (***R***=0) in the dry season and constant high rainfall (***R***=1) in the wet season.

With the rainfall data of a particular year and the survival function (eq. 10), we can now calculate each seedling’s survival probability (Fig. 2C), which is given by the product of its (age- and rain-dependent) survival probabilities of each week from germination time *d*_z_ until the end of the year:

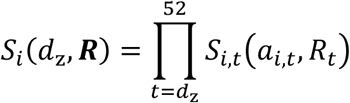

To determine whether a seedling survives its first year, we draw a random number from a uniform distribution [0,1]. If this random number is lower than the survival probability of the seedling, the seedling will survive, otherwise it is removed from the population. Likewise, we can calculate which of the adults will be allowed to reproduce (see below) and survive to the next year. We assume that weekly survival for plants of age *a*_*i*_ ≥ 27 weeks, is independent of age and rainfall and equal to *s*_ad_, such that yearly survival becomes *s*_ad52_ and expected lifespan (in weeks) is 1/(1-*s*_ad_).

#### Recruitment and density dependence

We assume that only *K* adult plants can live in the environment. At the end of the wet season, we determine which of the seedlings that germinated in this year will recruit into the adult population. We first calculate the number of available slots (*K* minus the number of surviving adults), and then fill the empty slots with seedlings. Recruitment success depends on seedling age, with older seedlings having a higher probability of recruiting. The relative recruitment probability of a seed is calculated as

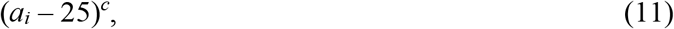

where parameter *c* indicates the strength of competitive asymmetry. For *c* = 0, recruitment is independent of age, while for high values of *c*, young seedlings will be outcompeted by older seedlings. Because seeds germinate at the latest in week 27, the youngest seedling is 26 weeks old at the time of recruitment. By subtracting 25 from the age of each seedling, the youngest seedling has a weight of 1, and all other seedlings have weights exceeding 1.

#### Reproduction, inheritance and mutation

At the end of the wet season, all adult plants (i.e. individuals older than 52 weeks, not the juveniles that were just recruited) reproduce asexually. Individual seed production is Poisson distributed with a mean fecundity of *γ*. Offspring inherit their parent’s gene values for determining dormancy duration (*d*_*i*_), but not their actual phenotype *d*_z_. Each locus has an independent probability *μ* of experiencing mutation. Mutational effects are of constant effect, such that the new gene value is drawn from a normal distribution with a mean equal to the previous value, and a standard deviation of *μ*_σ_. Mutations that would lead to negative values are absorbed at 0, ensuring that the new gene value is non-negative. After mutation, the values of all genes of an individual are normalized to sum to 1.

#### Data generation and analysis

Simulations run for 1.000.000 years. Results presented are based on the populations at the end of each simulation. As there is hardly variation among replicates, results are presented for only 3 independent replicate runs of each scenario. We implemented the model in C++; the code will be made available online. Simulations were run using GNU parallel (Tange 2021).

## Results

Despite the added ecological complexity and evolutionary realism of the individual-based simulation model, with its potential for a detailed flexible germination strategy, we overall find that these results correspond well with those of the analytical model (section 1). Firstly, despite the high potential for finely-tuned germination strategies, our simulations showed remarkably little variation both among replicated simulations and within populations, in agreement with our prediction that any ESS is convergence stable and so no evolutionary branching should occur. Secondly, our simulations highlight the mechanisms modulating the (rainfall-dependent) benefits and costs of early relative to late seeds – namely the strength of age-dependent competitive asymmetries for recruitment and juvenile dry-season survival – and corroborate the finding from our analytical model that mixed germination strategies only arise at a certain balance point where the relative benefits of early seeds are neither too large nor too small (Figs. 3, 4).

**Figure 3:**
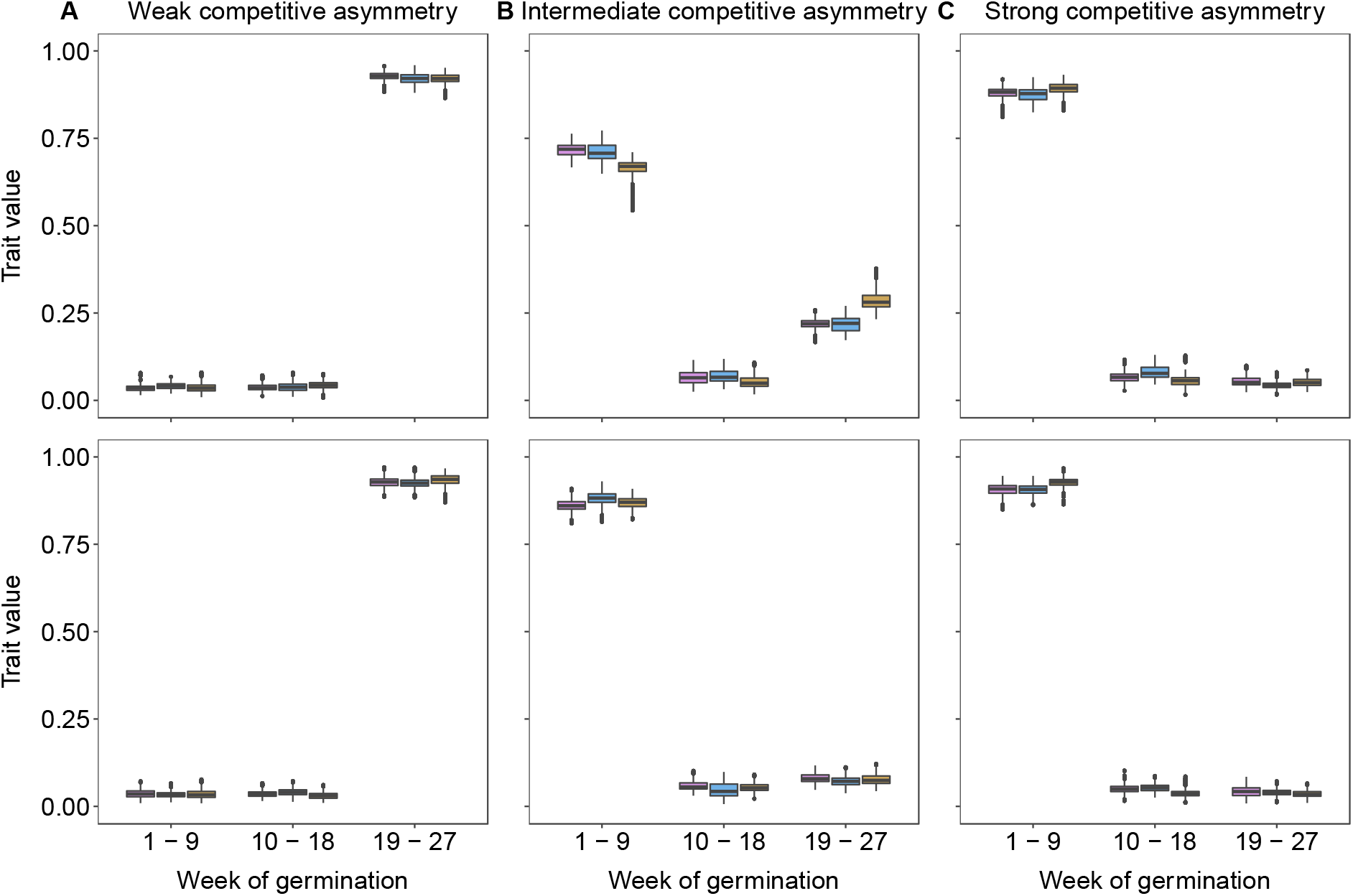
Evolved trait values (weekly germination probabilities) in three independent simulations (box colors) under increasing levels of competitive advantage to older seeds (‘competitive asymmetries’, columns). For clarity, results are shown summing the germination probabilities of 9 weeks together. Box plots show median (horizontal line), range of 50 % of variation (box limits), range of 95 % of variation (vertical lines) and any outliers (black points). Top row: Coarse-grained environment (all seedlings experience same rainfall patterns within years). Bottom row: Fine-grained environment (all seedlings experience different rainfall patterns within years). Strength of competitive asymmetry is determined by parameter *c*, which takes the values of 0.5 (left column), 1 (middle column) or 1.2 (right column). Other parameters as in table 1.

**Figure 4:**
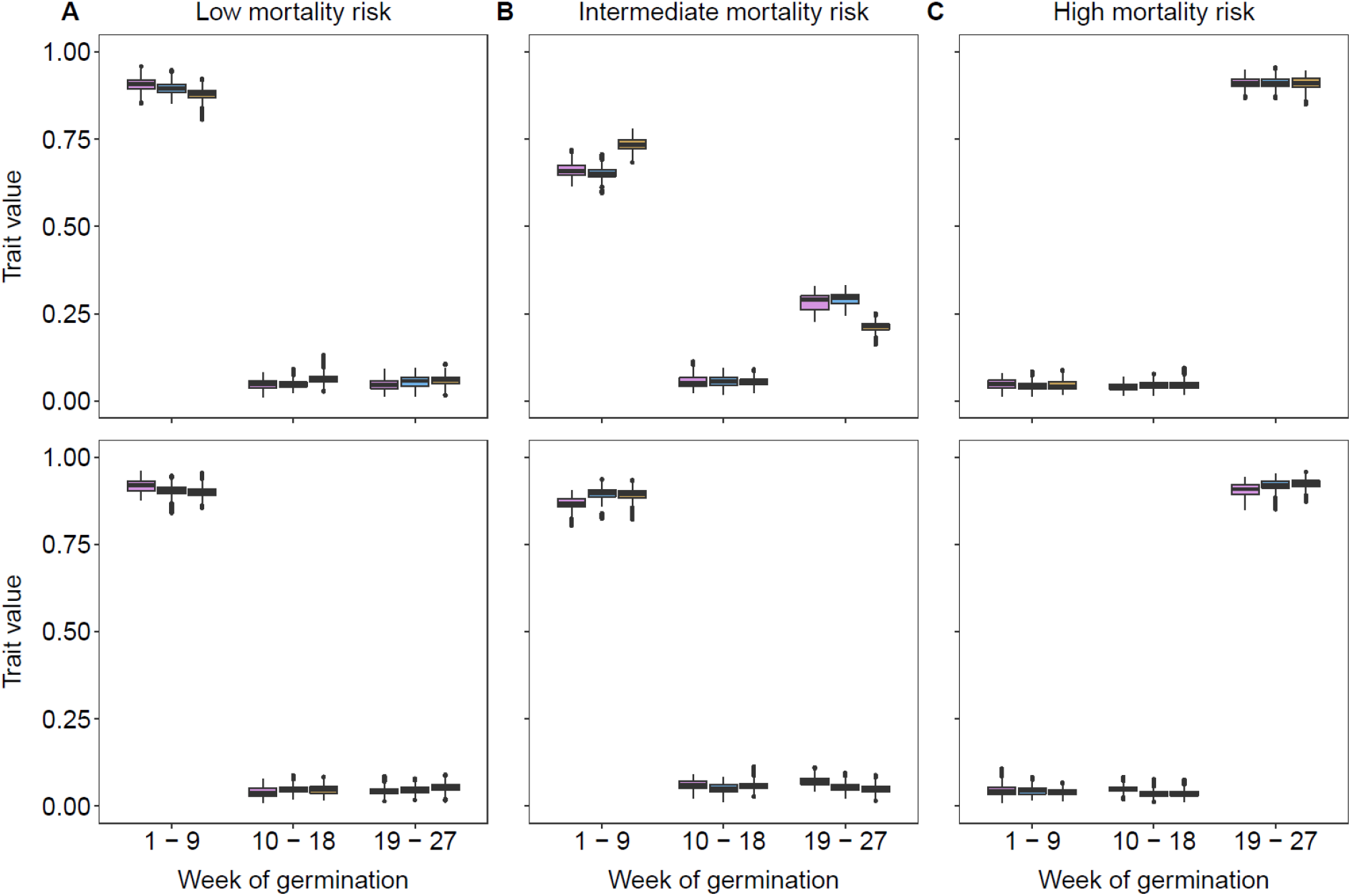
Evolved trait values (weekly germination probabilities) in three independent simulations (box colors) under increasing mortality risks (columns). For clarity, results are shown summing the germination probabilities of 9 weeks together. Box plots show median (horizontal line), range of 50 % of variation (box limits), range of 95 % of variation (vertical lines) and any outliers (black points). Top row: Coarse-grained environment (all seedlings experience same rainfall patterns within years). Bottom row: Fine-grained environment (all seedlings experience different rainfall patterns within years). Mortality risk is modulated by adjusting the *a*_inf_ parameter, which takes the values 4 (left column), 5 (middle column) and 7 (right column).

Additionally, expanding on the findings from the analytical model, our simulations demonstrate that this within-individual dormancy variation evolves as an adaptation lowering genotype-level fitness variance, confirming that such strategies do indeed represent bet-hedging strategies (compare top vs. bottom rows of Figs. 3, 4). By comparing results obtained in temporally coarse-grained environments (all individuals experience the same rainfall patterns in a given year, resulting in high within-year fitness correlations and high among-year fitness variation) with those obtained in fine-grained environments (rainfall patterns are assigned independently for all individuals in a given year, resulting in low within-year fitness correlations and low among-year fitness variation), we observe that any spreading of germination disappears in fine-grained environments. Thus, the evolution of within-individual seed size variation in perennials can here be attributed to the interaction between age-dependent survival, competition for recruitment, and bet-hedging. This result also importantly highlights that bet-hedging in germination strategies is not driven *per se* by weekly rainfall variation within years, *V*, but also by the spatial scale of rainfall variation, which affects correlations in environmental conditions experienced among individuals within populations.

When competitive asymmetries are weak, i.e. competition for recruitment only depends weakly on seedling age, obligate dormancy evolves causing germination at the beginning of the wet season (top row of Fig. 3a and S1a). This holds even under higher rainfall levels, for which dry-season germination and seedling survival may often be possible, because there are no competitive benefits to early germination and so even small risks are not worth taking. Conversely, when competitive ability depends strongly on seedling age, minimal dormancy periods evolve and seeds germinate as soon as possible (top row of Fig. 3c and S1c). These seedlings then need to survive the entire risky dry-season, but this pays off as long as the dry season is not too malign, because later-germinating seeds have little chance of recruiting if there are even a few surviving early seeds. Accordingly, levels of within-individual variation in dormancy period are low in both these scenarios. Bet-hedging has no effect on the evolution of dormancy in this case, because there is no variance in expected fitness nor fitness correlations among related individuals (top and bottom rows are qualitatively identical in Fig. 3a,c and S1a,c). Such fixed strategies and absence of evolutionary conflicts of interest in the short and long term are also predicted by our analytical model (eq. 8, no mixed strategies (0<*f*<1) can arise if the benefits *b* of germinating early do not increase sufficiently despite excess rain).

Similarly, only intermediate mortality risks select for mixed strategies. Because seedling mortality strongly affects the risks of early germination for a given rainfall level, any variation in germination timing occurs at intermediate levels of mortality risk in the dry season (Fig. 4). In the scenario shown in Fig. 4 we adjust the parameter *a*_inf_, the inflection point of the survival function, i.e. the age at which survival rises steepest, such that higher (later) *a*_inf_ entails higher overall mortality (see Fig. 2). We find that low mortality (left column in Fig. 4) always selects for germinating as soon as possible (because of age-dependent competitive asymmetries). On the other hand, very high mortality (due to high *a*_inf_; right column in Fig. 4) always selects for germinating as close as possible to the start of the wet season. Only intermediate mortality risks (Figs. 4 & S2, middle column) favor variable dormancy periods. Again, this general result corresponds well with our analytical model (Fig. 1).

A final observation in this model is that, in contrast with predictions from previous bet-hedging theory, shorter lifespans may select for less spreading of germination (Fig. 5, weekly results shown in Fig. S3). This effect arises because of our implementation of density-dependence, where lower adult survival means that more ‘slots’ are opened up every year for seedlings to recruit into. This decreases the intensity of age-dependent competition, leading to weaker selection for early germination the lower the adult survival (Fig. 5A shows results for high adult survival, versus Fig. 5B for low adult survival).

**Figure 5:**
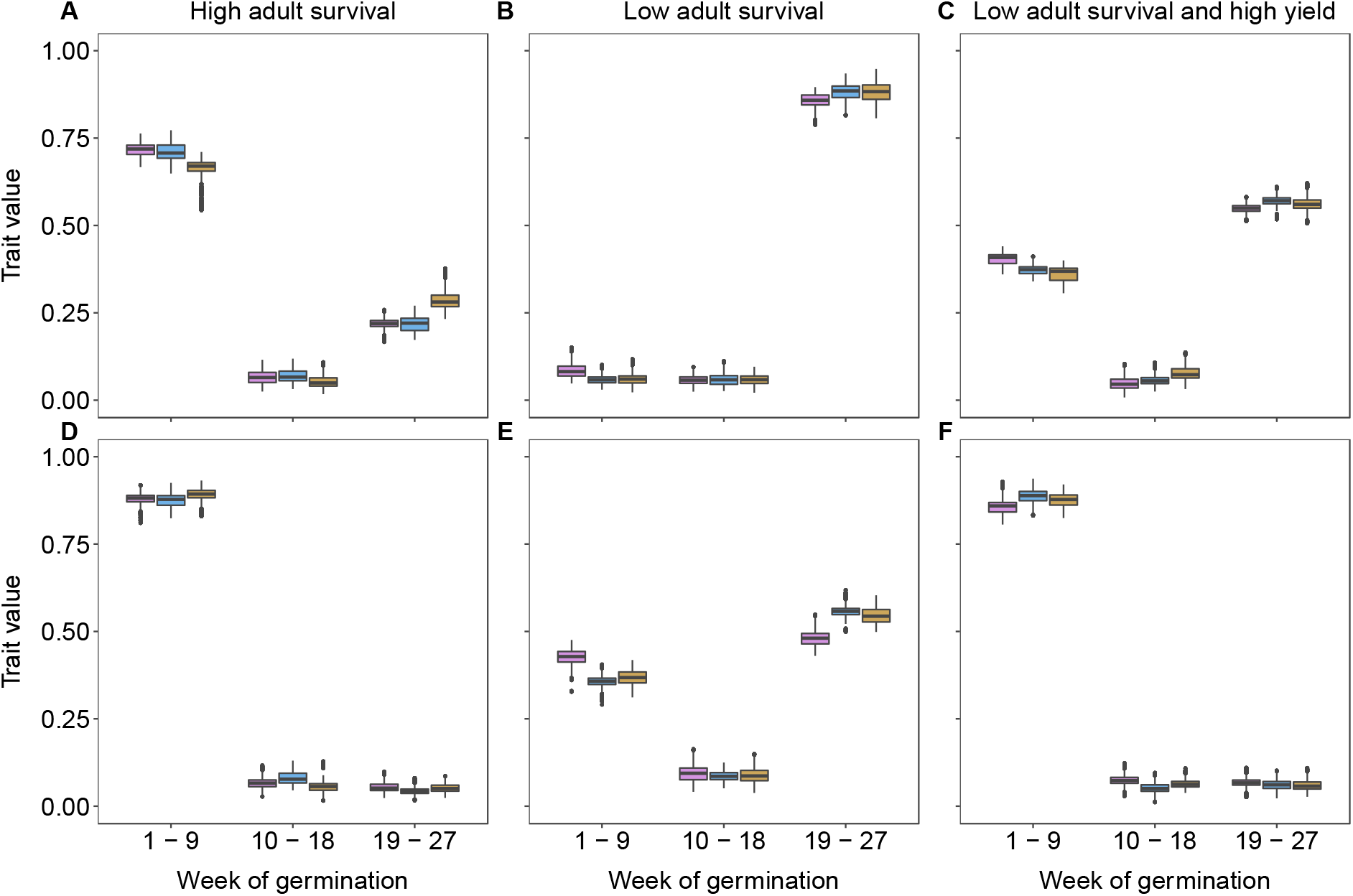
Evolved trait values (weekly germination probabilities) in three independent simulations (box colors) under different strengths of density dependence (columns) and levels of competitive asymmetry (rows). For clarity, results are shown summing the germination probabilities of 9 weeks together. Box plots show median (horizontal line), range of 50 % of variation (box limits), range of 95 % of variation (vertical lines) and any outliers (black points). Top row: Intermediate competitive asymmetry (*c* = 1), bottom row: strong competitive asymmetry (*c* = 1.2). Density-dependence is modulated by adjusting the adult survival rate *s*_*a*_ (0.997 in the left panels, 0.993 in the middle and right panels) and the average seed production (50 in the left and middle panels, 130 in the right panels).

In situations where competition is strongly asymmetric (high value of *c*, bottom row of Fig. 5), lowering adult survival may select for more spreading of germination (Fig. 5D, shows results for high adult survival, versus Fig 5E for low adult survival). The reduction in density-dependence now intensifies selection for late germination, resulting in the evolution of more spread in germination timing. In the analytical model, adult survival affects the speed of evolution only, and not the evolutionary endpoint (equation 4). This discrepancy with the individual-based model is caused by the assumption in the analytical model that the strength of density-dependence depends on the relative benefit of germinating early and the frequency of this strategy. In the individual-based model, however, the strength of density-dependence is also affected by the number of free spots and competing seedlings. If we reduce adult survival and increase the average seed production at the same time, such that the strength of density-dependence (expected number of seeds competing per open slot) remains the same, we no longer see an effect of adult survival on the resulting strategies (Fig 5C and 5F show the same pattern of uni- or bimodal germination as do Fig 5A and 5D).

As for Figs. 3 & 4, removing any effects of bet-hedging by implementing fine-grained rainfall patterns (and thus removing environmentally-induced fitness correlations among related individuals) leads to populations with only a single, fixed strategy in all scenarios depicted in Fig. 5 (results not shown). Thus, within-individual variation in germination timing can arise due to interactions between life-history traits, bet-hedging and intensity of intraspecific competition.

## Discussion

### Modelling results and insights

Our two complementary modelling approaches have shown how stochastic seasonal environments can select for within-individual variation in germination behavior through interactions between bet-hedging, life-history traits, and competitive asymmetries due to age-dependent seedling recruitment. First, our analytical model considering discrete variation (early-vs. late-germinating seeds) reveals ample scope for mixed germination strategies in long-lived species. This model revealed that the key determinant of whether variable germination evolves is the shape of the function relating how all benefits and costs of early vs late seeds change depending on dry-season rainfall environmental conditions. Specifically, both the risks of early germination if conditions are too dry before the beginning of the wet season, and the benefits provided by competitive advantages over later-germinating seedlings (who will be smaller at the time of competition for recruitment), are captured in such a function, which is shown to have to be increasing and accelerating around 0 (expected dry-season rainfall), but not too quickly, in order for a mixed strategy to evolve.

Inspired by these results, we next built an evolutionary simulation model of dormancy period, where we included both these benefits and costs to allow them to vary independently, as well as continuous within- and among-individual variation in seed dormancy. To allow unconstrained evolution of flexible germination strategies, we modelled the joint evolution of 27 unlinked genes representing weekly germination probabilities of offspring. Despite this flexibility, the results from our analytical model with only two discrete germination strategies were largely recovered: Mixed strategies emerged only when the potential benefits (competitive advantage) and costs (dry-season mortality) of earlier germination were neither too strong nor too weak, but were still observed for a large range of parameter values. Furthermore, intermediate dormancy durations emerged very rarely, with evolution rather favoring mixed parental strategies producing some early- and some late-germinating offspring. Through comparisons with evolutionary outcomes in fine-grained environments, which never exhibited such mixed strategies, we conclude that this represents a diversifying bet-hedging strategy evolving due to genotype-level fitness correlations. In coarse-grained environments, phenotypically variable offspring ensure that fitnesses of siblings become decoupled, thus ensuring low genotype-level variance in fitness (Starrfelt and Kokko 2012).

Our work demonstrates that explicit consideration of density-dependence is necessary for understanding the adaptive value of within-individual variation in offspring phenotype in long-lived organisms. Variation in offspring size within individuals is not only common in perennial plants, but also in many other iteroparous organisms including arthropods (Fox and Czesak 2000), birds (Amundsen and Slagsvold 1998), fish (Einum and Fleming 2004), and marine invertebrates (Marshall and Keough 2007). While among-population variation is often interpreted as adaptive, within-individual variation, on the other hand, is mostly viewed as maladaptive (Fox and Czesak 2000). This interpretation is possibly driven by the focus on optimality models (e.g. Smith and Fretwell 1974; Einum and Fleming 2004), which by definition predict a single offspring size to maximize fitness (Metz et al. 2008). Approaches that take frequency-dependent interactions into account, however, show that within-individual variation in offspring size can be adaptive, even in the absence of environmental unpredictability (e.g. Geritz 1995). Our work shows that such an ecological approach is also necessary to understand evolution in unpredictable environments.

### Germination strategies in seasonal environments

Our models were inspired by the wet-dry seasonal dynamics characterizing large parts of the tropics (Feng et al. 2013). Yearly patterns of seedling emergence such as those described by Garwood (1983) for Barro Colorado Island in Panamá suggest that germination during the early part of the rainy season predominates, although there is considerable variation (see also Escobar et al. 2018). Garwood’s study also hinted at an important role of asymmetric competition in that seedling emergence in highly competitive light gaps tended to occur earlier than did emergence in less competitive understory habitats. Our models provide a mechanistic explanation for these patterns, and also suggest that ongoing changes in the predictability and temporal variation in precipitation patterns in the tropics may select for altered germination strategies in tropical plants (Feng et al. 2013).

The adaptive challenges faced by plants of seasonal tropical forests are in some ways analogous to those of arctic-alpine plants facing the problem of ‘false springs’ followed by freezing events, which is thought to select for dormancy or other mechanisms to avoid detrimental fitness effects of premature germination (Schwienbacher et al. 2011; Mondoni et al. 2012). Arguing that within-individual variation in germination timing could be beneficial as a bet-hedging trait, Simons & Johnston (1997) used a model of such a system to illustrate their argument that developmental instability could be adaptive in annuals. During a time when developmental instability was largely seen as maladaptive (c.f. the debate surrounding fluctuating asymmetry), substantial work on bet-hedging strategies has largely resolved this controversy (Zhang and Hill 2005; Devaux and Lande 2010; Scheiner 2014; Tufto 2015), and the dynamics considered by Simons & Johnson (1997) are now a canonical example of bet-hedging in seasonal phenology.

A second source of inspiration for our models was the extensive variation in seed size often observed in angiosperms (Michaels et al. 1988), and the many studies reporting relationships between seed size and germination behavior (e.g. Simons and Johnston 2000; Norden et al. 2009; Martins et al. 2019). Although seed size is sometimes thought about as a highly canalized trait (because there should be an optimal balance between size and number of offspring; Smith and Fretwell 1974), both categorical (seed heteromorphism) and continuous variation in seed size is common (e.g. Susko and Lovett-Doust 2000). In a recent phylogenetic analysis, Scholl et al. (2020) analyzed the occurrence of seed heteromorphism in a large sample of the flora of southwestern North America, and detected weak associations between seed heteromorphism and several measures of environmental predictability (aridity and precipitation seasonality). While the tendency for seed heteromorphism to be more likely in more unpredictable environments is consistent with our simulation model, these authors explicitly excluded species with continuous variation in seed size from their analyses. Although our simulation model suggests that seed heteromorphism may be a more likely evolutionary outcome for dealing with environmental variability than continuous seed size variation, there is no reason why continuous seed size variation could not also represent a bet-hedging strategy in unpredictable environments (c.f. Simons & Johnston’s (1997) suggestion of developmental instability causing adaptive variation in germination timing). Interestingly, Scholl et al. (2020) failed to detect a correlation between seed heteromorphism and the annual life cycle, contrary to their expectations from previous bet-hedging theory. Although seed heteromorphism was more common in annuals than in perennials, they attributed the lack of a detectable effect to the small overall proportion of seed-heteromorphic species. However, the common occurrence of seed heteromorphism also among perennials can further be seen as an indication that the processes interacting with bet-hedging in our model to produce variation in germination phenology may indeed be common and important in natural systems.

### Empirical guidelines and predictions

Our models yield several testable predictions about the relationship between germination behavior, seed size, and environmental predictability. The simplest prediction arising from our results is that within-individual variability in germination behavior and associated traits (e.g. seed size) can be adaptive, and possibly more so in more unpredictable environments. However, we also find that the distinctions between fine-vs. coarse-grained environmental variation is important in driving germination patterns. This implies that caution is needed when interpreting measurements of environmental variation and using them to predict ecological gradients of germination strategies. For example, the measurement of rainfall variation used in Scholl et al. (2020) was based on among-year variation (coefficient of variation of annual rainfall measurements) and the smallest spatial grid used was 25 km^2^, which is too large to capture whether a given genotype may experience environmental variation as fine- or coarse-grained. Thus, neither of these metrics provide information about the different outcomes observed in our models, which might explain the lack of a relationship between rainfall variation and the occurrence of seed heteromorphism. Another reason highlighted by both our models could be that mixed germination strategies as adaptations to unpredictable environments may only evolve under certain conditions of how fitness costs and benefits vary among morphs across environments.

Recent studies of variation in seed size and germination behavior in the tropical vine *Dalechampia scandens* have revealed predictable covariation between measures of environmental predictability and germination behavior. Martins et al. (2019) found that seeds from populations occupying more seasonal environments required longer periods of after-ripening (duration of storage required before germination occurs in response to favorable wet conditions) before germinating, and that within each population, smaller seeds required shorter periods of after-ripening. In a follow-up study, Pélabon et al. (2021) considered variation in seed size within individuals and detected a positive relationship between environmental stochasticity and variation in seed size. Environmental unpredictability as defined by Pélabon et al. more closely resembles our parameter *V* in describing stochastic variation in rainfall patterns, and our model thus provides mechanistic support for the hypothesis that the patterns of seed size variation and germination behavior in *Dalechampia* represents a bet-hedging strategy in unpredictable environments.

While observational studies of germination timing and seed size across ecological gradients may continue to yield insights into variation in these traits, experimental studies would be highly valuable. Seed-sowing experiments with seed families of known descent would specifically allow separating environmental and genetic causes of variation, and lead to insights into the evolutionary potential of germination behavior (e.g. Simons & Johnson 2006). However, separating the presence of two distinct strategies from continuous variation may continue to prove difficult, especially if the difference in mean behavior between strategies is small, and variation around these means is large.

### Limitations and caveats of the model

Our model results are broadly consistent with empirical patterns of seed size and germination behavior, yet some caution is needed when interpreting the model output. In particular, the specific genetic architecture we use (one freely evolving locus for each weekly germination probability) is admittedly highly unrealistic. Although we hope that this flexibility has allowed us to observe outcomes that explore all parts of unconstrained strategy space, some potentially beneficial combinations of germination probabilities may have been difficult to arrive at. For example, by linearly standardizing all loci of weekly germination probabilities to sum to 1, loci with already large values could make the effect of potentially beneficial mutations at other loci increasing the germination probability in other weeks very small.

Next, while this breach in realism aimed to limit bias and increase the robustness of our evolutionary interpretation, other limitations may in fact introduce biases to our conclusions. For example, it is difficult to say how our choice of asexual inheritance might have affected the ability to evolve discrete vs. continuous strategies, or indeed the range of parameter space favoring variable germination, and so including sexual reproduction could provide an interesting next step for expanding our modelling work. Plant mating systems are highly variable, and near-complete selfing is uncommon compared to mixed and predominantly outcrossing systems (Moeller et al. 2017). If germination timing is indeed highly polygenic, outcrossing among individuals producing predominantly early- and predominantly late-germinating seeds might allow mixed strategies to evolve more easily than in our model, where such mixed strategies instead are dependent on the right mutations occurring sequentially. In addition to dormancy, germination timing is also driven by variation in flowering and fruiting phenology as well as the duration of fruit maturation (Escobar et al. 2018). Although we chose to keep these processes fixed in our model, variation in flowering time additionally leads to non-random mating and is therefore an interesting trait to add if future studies consider sexual reproduction.

Modelling multilocus evolution in the absence of any constraints, as we do here, may in itself introduce biases if evolution of optimal combinations is difficult to attain due to genetic constrains, covariation with other traits, or trade-offs. Importantly, we did not incorporate a trade-off between offspring quality and dormancy period, which could be expected if we assume that seed size is the mechanism by which variation in germination strategy is achieved (Stearns 1989). Furthermore, we did not include a cost to seed size variation (or canalization, see Zhang and Hill 2005). However, while such additional assumptions might affect our results (for example by narrowing the parameter space in which variable strategies are selected for if adding a cost to variation, and widening it if adding a cost to canalization), they also reduce generality and do not subtract from the general point. Variation in germination strategies may arise from other sources than simply seed size variation resulting from per-offspring parental investment (Simons and Johnston 2006; Baskin and Baskin 2014) and as with costs of variation vs. canalization, we note that the pattern may also go either way with regards to the seed size vs. germination strategy (offspring quantity vs quality) trade-off: In many species, larger seeds germinate earlier (Biere 1991; Simons and Johnston 2000; Pélabon et al. 2005), but the opposite pattern is also observed (Susko and Lovett-Doust 2000), and among species time to germination is usually longer for larger seeds (Norden et al. 2009; Harel et al. 2011). Thus, while adding some trade-off between seed set and germination strategy might increase realism with respect to certain systems, generality would again necessarily be lowered relative to our present model without such energetic constraints.

## Conclusions

This study has demonstrated novel mechanisms by which variation in germination strategies in perennial plants can evolve, shedding light on the prevalence of seed-size variation seen in perennials as well as annuals. Seed size was historically considered a highly canalized trait, to the point that the weight measurement ‘carat’ was based on the remarkably invariant seeds of the carob tree (*Ceratonia siliqua*). However, it has since been discovered that substantial variation in seed size is widespread, even in carob seeds (Turnbull et al. 2006), and its universality begs an explanation. Bet-hedging in desert annuals has previously been the best-studied empirical and theoretical example of variable germination strategies, but there are arguably more perennial species occupying seasonal tropical or temperate environments than there are desert annuals. Our modelling results, where bet-hedging in perennials interacts with unpredictable seasonal variation and competitive asymmetries to produce within-individual variation in germination strategies, therefore provide a much-needed extension to existing theory on this topic. A better understanding of competitive dynamics in stochastic seasonal environments can help improve the predictability of species and community responses to ongoing changes in climate patterns and other anthropogenic challenges.

## Acknowledgments

The authors thank Hanna Kokko for helpful discussions, and an anonymous reviewer for a great contribution to the analytical model. HtB and TRH were supported by the Swiss National Science Foundation, on grant no. 310030B_182836 awarded to Hanna Kokko. TRH was additionally supported by the Forschungskredit of the University of Zurich, grant no. FK-21-122 awarded to TRH, and the Research Council of Norway, grant no. 313570 awarded to Jane Reid.

## Author contributions

All authors initiated the study. HtB and TRH developed and analyzed the models. HtB ran the simulations. ØHO reviewed the empirical literature. TRH reviewed the theoretical literature and drafted the manuscript with input from all authors.

## Appendix A: Derivation of invasion fitness (eq. 2)

Following a standard adaptive dynamics approach (Geritz et al. 1998), we assume sequential appearance of rare mutations of small phenotypic effect in populations of residents fixed for the wild type, where each mutation is assumed to go to fixation (if its invasion fitness is positive) before the next mutation occurs. Mutants are assumed to have negligible effect on ecological dynamics, such that the number of mutants *N’* in a population of *K* residents can be derived. As in the main text, we designate respectively the resident and mutant germination strategies (fraction of seeds early) as *f* and *f’*, and probability of recruitment for an early seed is given by eq. 1. Then, the dynamics of number of mutants *N*’(*t* + 1) is given by the number of surviving mutant adults from the previous time step, *N*’(*t*)*s*_a_, plus any mutant seedling recruitment (eq. A1). As described in the main text, seedling recruitment is according to a lottery model where all seeds compete for a limited number of spaces in the adult population freed up after adult mortality; this number is (1 – *s*_a_)*K*. Assuming equal yields of *γ* seeds per mutant and wild type adult, the number of mutant seeds produced is *N’*(*t*) *γ*, and the number of resident seeds is *K γ*, and their mean (averaged over small and large seeds) recruitment probabilities are respectively *f’b* + 1 – *f’* and *fb* + 1 – *f*. The number of mutants in the next time step is given by

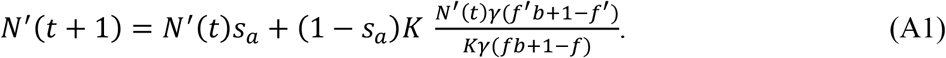

Cancelling away *K γ* and dividing by *N’*(*t*) gives the growth rate *λ* of the mutant population used in eq. 2:

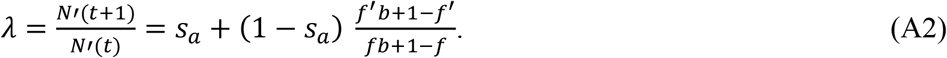

## Data availability statement

All code and output is available upon request and will be uploaded to Dryad upon journal acceptance.

## Supplementary material

**Figure S1:**
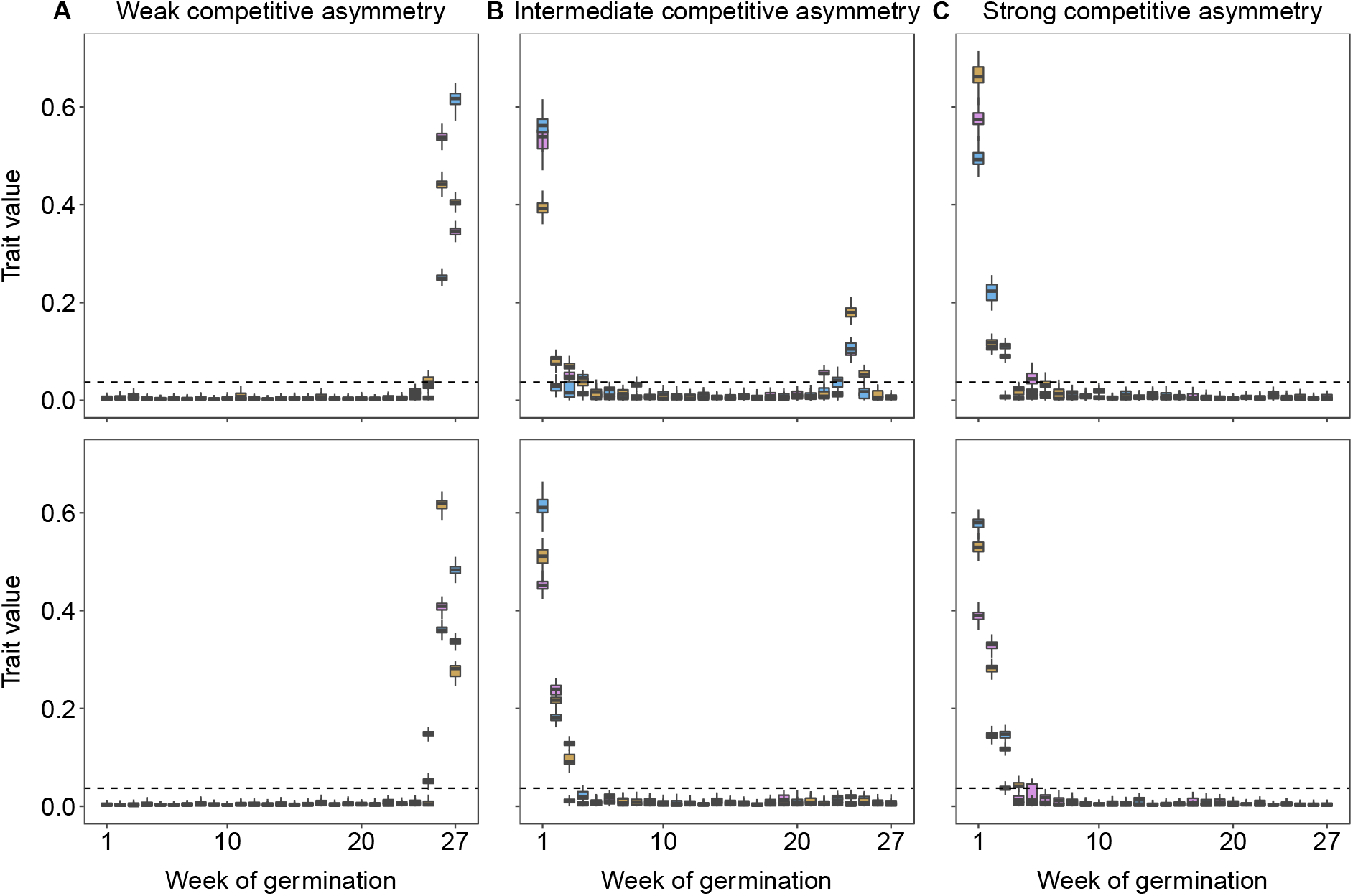
As Figure 3, except with each weekly germination probability plotted individually rather than grouped by 9 weeks at the time. The dashed horizontal line indicates the probability 1/27, which would be the expected evolved value in the absence of selection on germination week.

**Figure S2:**
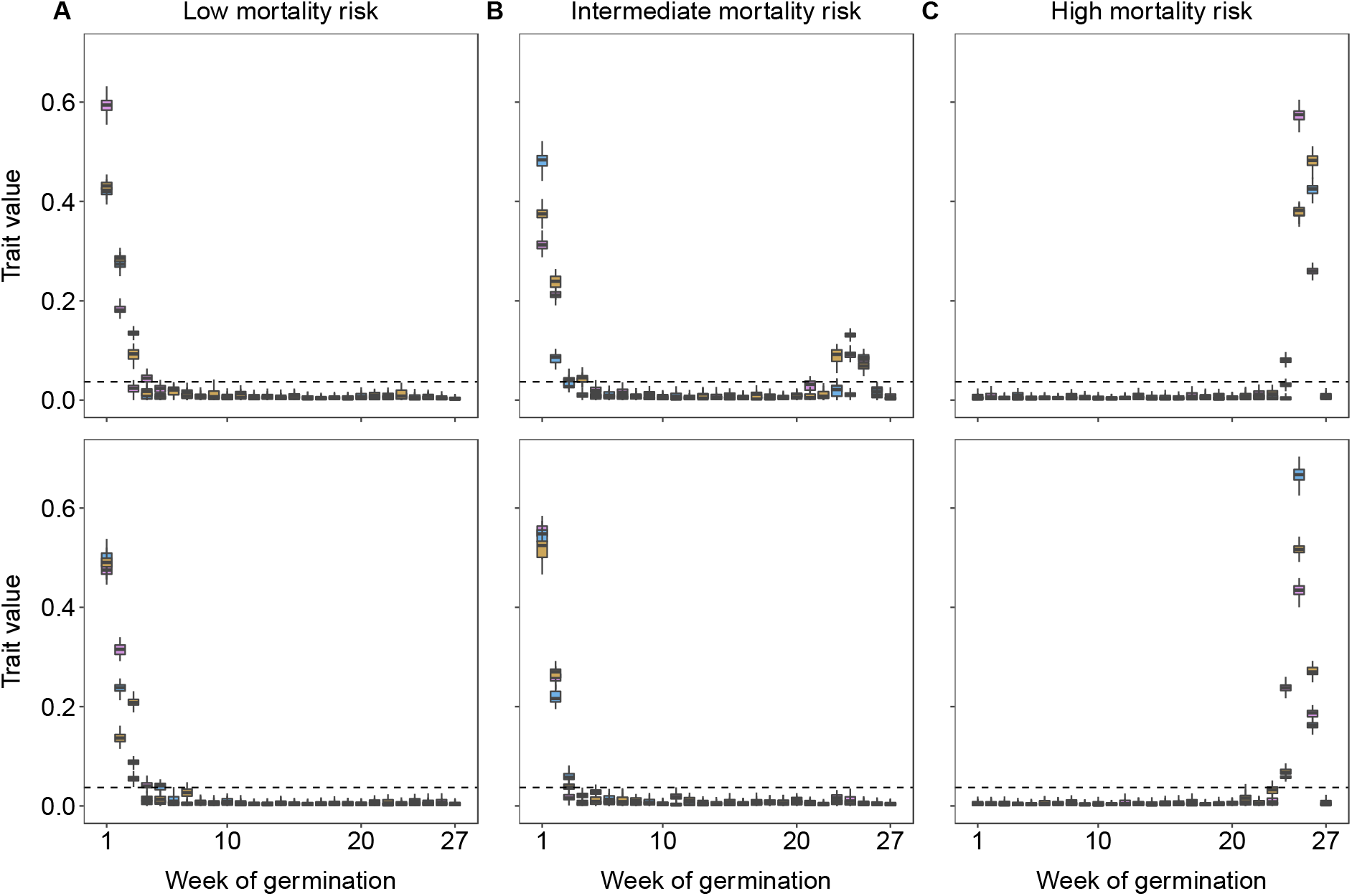
As Figure 4, except with each weekly germination probability plotted individually rather than grouped by 9 weeks at the time. The dashed horizontal line indicates the probability 1/27, which would be the expected evolved value in the absence of selection on germination week.

**Figure S3:**
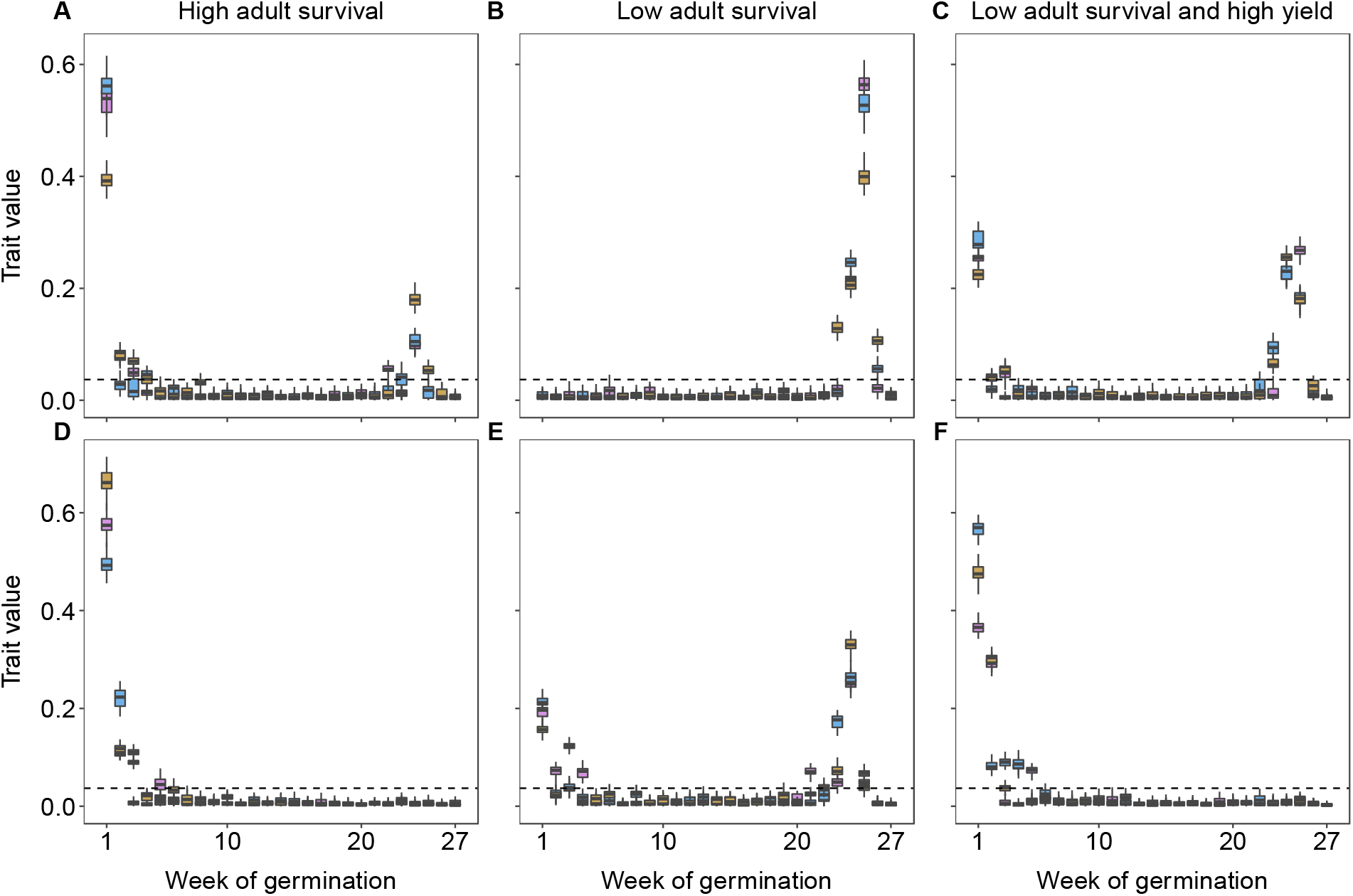
As Figure 5, except with each weekly germination probability plotted individually rather than grouped by 9 weeks at the time. The dashed horizontal line indicates the probability 1/27, which would be the expected evolved value in the absence of selection on germination week.

